# Stratifying variant deleteriousness and trait-modulating effect under human recent adaptation

**DOI:** 10.1101/2024.07.15.603534

**Authors:** Xutong Fan, Dandan Huang, Xinran Dong, Xianfu Yi, Jianhua Wang, Shijie Zhang, Xiaobao Dong, Xiaoqiong Gu, Miaoxin Li, Pak Chung Sham, Wenhao Zhou, Mulin Jun Li

**Affiliations:** Guangzhou Women and Children’s Medical Center, Guangzhou Medical University, Guangzhou, China; Department of Bioinformatics, The Province and Ministry Co-sponsored Collaborative Innovation Center for Medical Epigenetics, School of Basic Medical Sciences, Tianjin Medical University, Tianjin, China; Department of Pharmacology, Tianjin Key Laboratory of Inflammation Biology, School of Basic Medical Sciences, Tianjin Medical University, Tianjin, China; Center for Molecular Medicine, Children’s Hospital of Fudan University, Shanghai, China; Department of Genetics, School of Basic Medical Sciences, Tianjin Medical University, Tianjin, China; Program in Bioinformatics, Zhongshan School of Medicine, Sun Yat-Sen University, Guangzhou, China; State Key Laboratory of Brain and Cognitive Sciences, The University of Hong Kong, Hong Kong SAR, China

**Author notes:** Correspondence (W.H.Z), (M.J.L.). The authors contributed equally to this work (Xutong Fan, Dandan Huang).

## Abstract

Despite advances in annotating and interpreting human genetic variants, existing methods to distinguish deleterious/pathogenic from neutral variants still inadequately capture the nuanced impact of genetic variants on fitness and disease susceptibility. In this study, we introduced a new deep learning framework, the FIND model, by stratifying genetic variants into refined categories based on selection pressures and derived allele frequency. FIND demonstrated superior performance over existing genome-wide methods, delivering enhanced resolution in differentiating trait-modulating alleles from those that are pathogenic or neutral. Access to base-wise informative annotations has unveiled novel features that significantly enhance the model interpretability, and FIND has adeptly delineated evolutionary trends in human accelerated regions. Furthermore, applying FIND to the interpretation of clinical variants demonstrates its substantial potential in reclassifying variants of unknown significance. This work advances our understanding of the genetic underpinnings of human adaptation and disease, providing a new tool to explore the complexities of genetic contributions to health.

## Introduction

Although human adaptation is but a fragment within the extensive continuum of species evolution, deciphering the functional evolution of genetic variants influenced by recent natural selection is crucial (*1*). Specifically, understanding how these variants shape phenotypic traits, alter disease susceptibility, and even influence human fitness is essential for ultimately advancing health management and intervention (*2*). Nevertheless, current large-scale genomic sequencing or genome-wide association studies (GWASs) across different human subpopulations have only identified a limited number of true disease/trait-causal variants (*3, 4*) or positively selected loci (*5*). Emerging high-throughput techniques for genetic manipulation and single-cell sequencing across diverse human tissue and cell-derived models have profoundly enhanced the functional studies of a select group of critical human genetic loci and causal alleles (*6, 7*), however, our understanding of the function and pathogenicity of the vast majority of human-derived alleles remains incomplete.

Significant advancements have been made in the methods used to infer the roles and characteristics of alleles in human adaptation from the composition and allelic structures of modern human genomes (*8–13*), yet several persistent challenges remain. First, while genome-wide prioritization and unbiased comparison across different variant consequences have been widely adopted (*14*), these tools distinguishing deleterious/pathogenic from neutral variants currently fail to adequately address the fine-grained impact of genetic variations on organismal fitness, often overlooking the assessment and prioritization of a vast majority of alleles that are not unequivocally deleterious/pathogenic. Disease/trait-modulating alleles that moderately affect human fitness and are favored by recent adaptation can be captured through GWASs or by learning local genome structure from diverse human subpopulations (*3, 15, 16*). However, current methodologies still lack the capability to simultaneously model the negative and positive adaptation effects of human variations. Second, with the profound insights into the non-coding human genome provided by recent functional genomics studies, such as ENCODE (*17*) and EpiMap (*18*), there is a critical need for the integration of the latest large-scale variant functional annotations with novel interpretable computational strategies for more accurate prediction of non-coding alleles. Such efforts will bridge the gap between mere variant prioritization and their functional, and even clinical interpretation, in an evolutionary context.

In this study, we integrated a multi-class ensemble of genetic variants, extensive base-wise annotations, and a deep learning framework to develop the FIND model. This model enhances the resolution in stratifying deleterious alleles and trait-modulating alleles favored by recent human adaptation. Using comprehensive datasets, FIND demonstrated exceptional performance in predicting distinct trait-modulating alleles and effectively differentiated evolutionary trends in rapidly evolving human regions. We also reassessed a wide range of annotation features across various biological domains, identifying new features that enhance the model’s interpretability. FIND’s high sensitivity in classifying clinical pathogenic variants and its better control over false-positive rates, as evidenced by data from ClinVar (*19*), underscore its potential to transform our understanding of genetic contributions to human health and disease.

## Results

### Construction of FIND model based on fitness partition, large-scale features and deep learning

To enhance the resolution in recognizing the contributions of human genetic variants to fitness, we have expanded the conventional binary partitioning of all human genome variants into four nuanced categories, utilizing the full dimensions of natural selection and derived allele frequency (DAF) spectrum (**Fig. 1A**), including 1) Fixed/Nearly Fixed (**F**): we have identified that humans harbor derived alleles (inferred from the human-chimpanzee ancestral genome) with a frequency of at least 95%. Less than 5% of these alleles appear as nearly fixed polymorphisms in the 1000 Genomes Project (*20*) (1KGP) variant catalog (DAF ≥ 95%). Nearly all of these genetic events are fully fixed within the human lineage; 2) Intermediate Selection (**I**): this new category encompasses derived alleles that have adapted to diverse environments and lifestyles, or exhibit susceptibility or resistance to various pathogens and diseases. These alleles also demonstrate a trade-off between adaptive evolution and disease susceptibility, typically captured by the causal variants identified in thousands of GWASs for abundant human diseases/traits. The alleles, which modulate traits and have been favored by selection (specifically referred to in this study as variants with trait-modulating effects), display a wide range of DAFs throughout different stages of human evolution; 3) Neutral (**N**): we have pinpointed human-derived alleles located in evolutionarily non-conservative recombination hotspot regions. These alleles are unlikely to make a significant contribution to fitness, due to their genomic positioning; 4) Deleteriousness (**D**): as previously mentioned (*8*), *de novo* mutations were simulated using an empirical model of sequence evolution, which incorporates specific CpG dinucleotide rates and overall mutation rates. This model predicts a number of deleterious alleles that would be naturally purged through selection processes (see details in **Methods**).

**Fig. 1.**
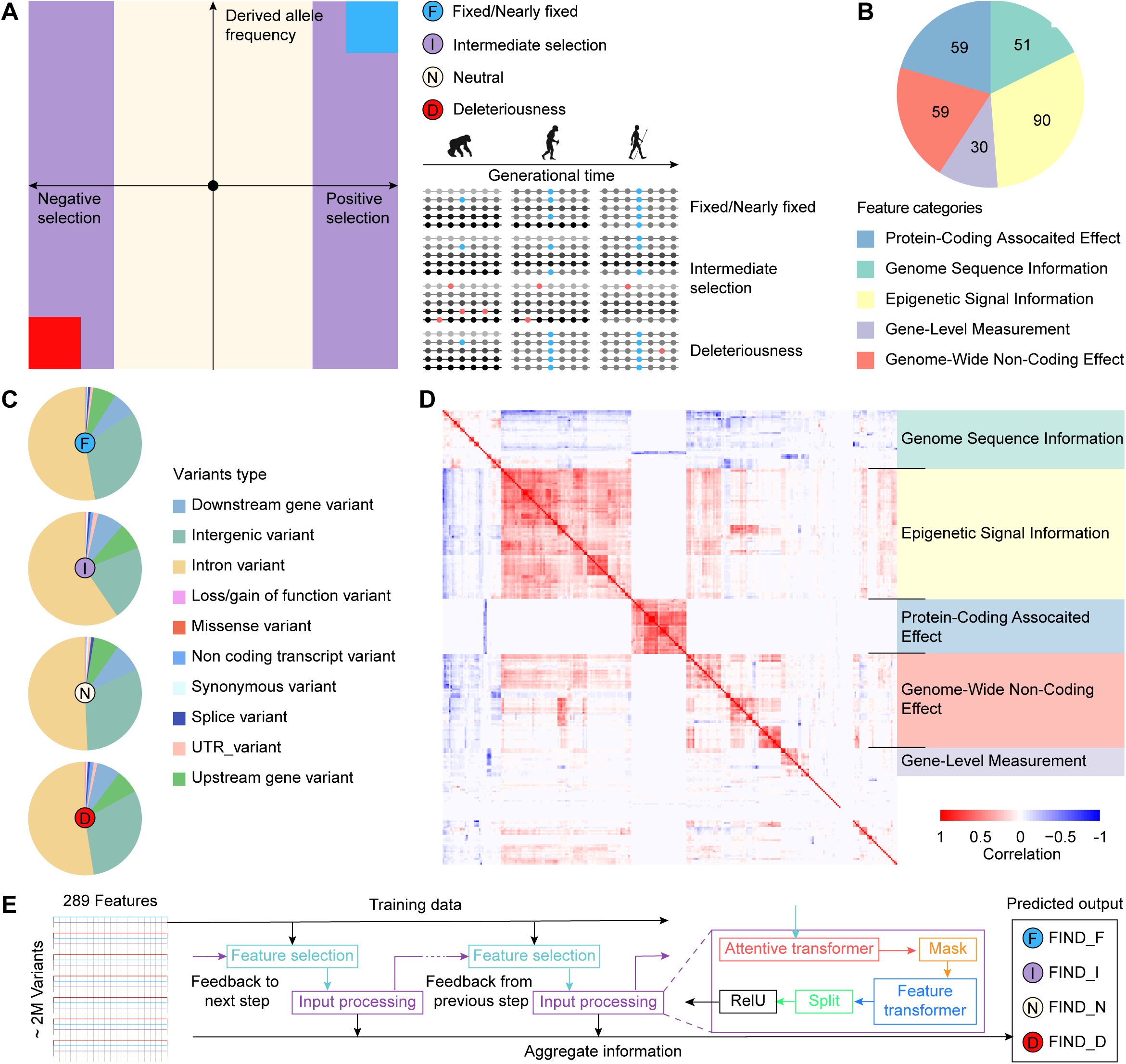
Development of FIND model integrating genetic fitness categorization, extensive features, and deep learning. **A**, Evolutionary basis for FIND models of variant fitness categories. Functional genetic architecture of variants from a spectrum of high derived allele frequency (DAF) driven by positive selection pressure (tend to fix in the human genome) to low DAF which is dominated by negative selection pressure (tend to disappear in the human genome), and neutral variants were random change in genome by both selection strength. It is simply explained in the timeline of human evolution: each dotted strand represents a haplotype. Alleles frequency that increases or decreases over time due to selection are represented as dots in light blue (advantage variants), red (deleterious or *de novo* variants), and shades of grey (linked variation to causal allele, neutral variants). We divide genome variants into four categories: Fixed/Nearly Fixed variants (**F**, blue), Intermediate variants (**I**, purple), Neutral variants (**N**, milk-white), and Deleterious variants (**D**, red). **B**, Pie chart showing training features composition, mainly divided into five categories. **C**, Variants consequence proportion of training datasets four categories in the pie chart. Variant consequences are obtained from the VEP. **D**, Heatmap of 289 features. Pearson correlation matrix calculated based on the training dataset. Positive correlations are displayed in red and negative correlations in blue color. **E**, Schematic view of FIND model, and FIND outputs the probability that the variant is classified into four categories: FIND_F, FIND_I, FIND_N and FIND_D.

To comprehensively capture the evolutionary patterns and adaptive selection mechanisms of human variations, particularly those in non-coding regions, we systematically computed and integrated 289 base-wise variant annotation features across the entire genome. These are stored in a terabyte (TB)-scale compressed file, with a significant percentage achieving allele-specific resolution (**Table S1**). Broadly, these annotation features can be categorized into five primary groups: 1) Genome Sequence Information (51): this category primarily consolidates genomic sequence characteristics and variant type statistics, providing a foundational layer of genetic context; 2) Epigenetic Signal Information (90): features in this category include aggregated scores for tissue-and cell-specific chromatin states, histone modifications, and transcription factor binding, sourced from comprehensive databases such as ENCODE, Epigenomics Roadmap (*21*), and EpiMap; 3) Protein-Coding Associated Effect (59): this group encompasses functional and deleterious impact scores on proteins, drawn from databases like dbNSFP (*22*), as well as scores predicting effects on alternative splicing; 4) Genome-Wide Non-Coding Effect (59): we have incorporated scores for evolutionary conservation (using metrics such as PhyloP (*23*), PhastCons (*24*), and GERP (*25*)) and many non-coding predictive scores from regBase (*14*), along with other genome-wide variant impact metrics, to assess the biological significance of non-coding alleles; 5) Gene-Level Measurement (30): multiple gene-level metrics have also been integrated and computed from various perspectives, including gene essentiality features, three-dimensional (3D) genome and target gene characteristics, and gene expression-related features (**Fig. 1B**, see details in **Methods**).

We constructed a training dataset comprising approximately 2 million variants, ensuring a balanced representation across four distinct categories, and annotated each variant with 289 allele-specific features. The distribution of variant consequences across these categories aligns with the actual genomic proportions of different variant types, which consistently showed a predominance of intronic and intergenic variants (**Fig. 1C**, **Table S2**), indicating that our sampling of training data was unbiased. Moreover, a genome-wide analysis of the correlation structure among the 289 features revealed that attributes within the same major category tended to cluster together, suggesting significant collinearity among some features, which necessitates careful consideration in subsequent model development to refine key features (**Fig. 1D**, **Table S3**). We employed the Attentive Interpretable Tabular Learning (TabNet) (*26*), a deep learning framework, to train multi-class classification of each variant based on its allele-specific features, called FIND. Unlike traditional machine learning frameworks such as ensemble learning, FIND leverages TabNet’s sequential attention mechanisms to selectively concentrate on the most pertinent features at each decision step. This method not only bolsters the desired biological interpretability but also enhances learning efficiency by focusing the model’s capacity on the most critical features (**Fig. 1E**).

### FIND accurately classifies genome-wide pathogenic variants

We initiated our analysis by comparing the performance of the FIND model trained using the TabNet algorithm with other commonly-used machine learning techniques. We observed that ensemble learning methods, like AdaBoost, outperformed traditional classifiers such as logistic regression (**Fig. 2A**, **Fig. S1A-B** and **Table S4**). Notably, the models trained using the TabNet algorithm exhibited the most superior prediction performance, demonstrating improvements of approximately 6.6%–17.2% in the area under the receiver operating characteristic curve (AUROC). Ten-fold cross-validation highlighted FIND’s exceptional multi-class prediction capabilities (average AUC = 0.970, **Fig. 2A**, **Fig S2B-E**; average AUPR = 0.926, **Fig. S2A, Fig S2F-I**). An ensemble average of these models was then applied to score all possible single nucleotide variants (SNVs), totaling over 8.6 billion, across the human reference genome. The raw scores were converted into Phred-like scores (PHRED scores) for more convenient comparison (*27*). By analyzing the distribution of PHRED scores across all possible SNVs and their specific functional consequences, we discerned distinct patterns in the distribution of FIND prediction scores among four outcome categories (**Fig. 2B**). Particularly, the FIND_D category displayed a unique configuration, where loss/gain of function variants (mean PHRED score = 9.716), missense variants (mean PHRED score = 8.161), and splice variants (mean PHRED score = 4.108) were highly prioritized and scored, aligning well with previous findings (*8*). Within the FIND_I category, although the proportion of above three variant types also increased with rising PHRED scores, they did not reach the extreme predominance observed in FIND_D. In contrast, some non-coding variants (such as those in 5′ and 3′ untranslated regions, non-coding transcript variants, and upstream gene variants) also occupy a modest proportion within the highly scored FIND_I category, suggesting that genetic variants with trait-modulating effects are also undergoing natural selection in the vast non-coding regions of the genome. Furthermore, in the FIND_F category, the proportion of non-coding and splice variants in the high PHRED score intervals increased, yet the proportion of loss/gain of function and missense variants remained low, indicating that many variants that do not directly disrupt protein-coding sequences may be more likely to exhibit advantageous phenotypes, being positively selected and fixed in recent human adaptation (*28, 29*). The distribution of variant types in the FIND_N category showed no significant correlation with PHRED scores, maintaining a relatively stable proportion across different intervals, implying that the model accurately captured the correct genomic neutral sites.

**Fig. 2.**
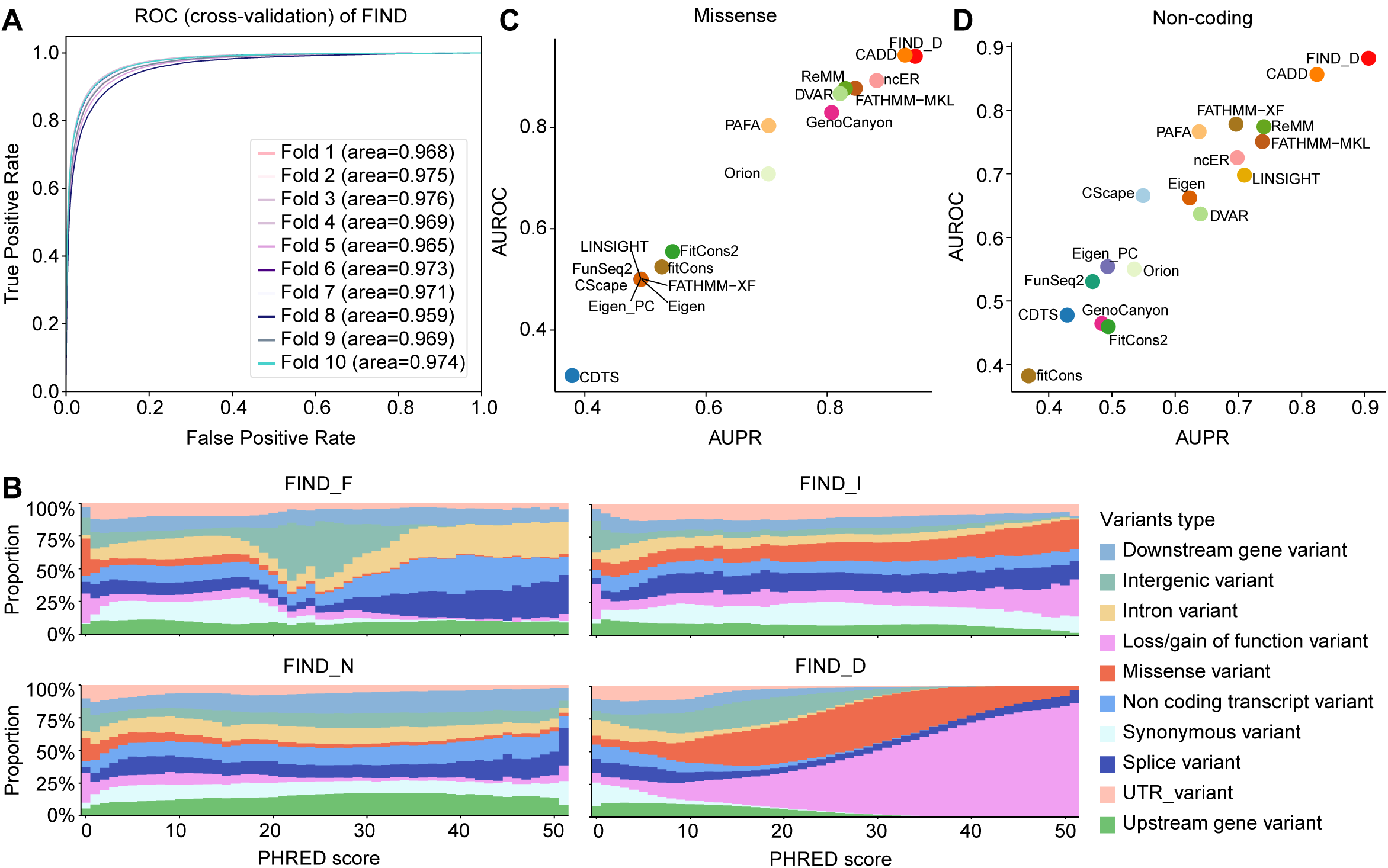
Assessment of FIND model’s accuracy in classifying pathogenic genome-wide variants. **A**, Receiver operating characteristic curve (ROC) and area under the receiver operating characteristics curve (AUROC) for FIND model ten-fold cross-validation evaluation (micro score was calculated because of four-classification task). **B**, Relationship of FIND four categories PHRED scores and genome-wide variants consequences, which shows the proportion of variants with a specific consequence in the continuous FIND PHRED score bin after normalizing by the total number of variants observed in that consequence category. **C**, AUROC and area under precision-recall curve (AUPR) of 17 tools, and FIND_D testing by missense variant dataset (19,772 pathogenic missense variants from ClinVar and 20,405 benign missense variants from gnomAD). **D**, AUROC and AUPR of 17 tools, and FIND_D testing by non-coding variant dataset (44 pathogenic regulatory variants from two recent publications and 58 benign non-coding variants from ClinVar).

To evaluate the performance of FIND in predicting well-documented pathogenic variants, we systematically curated and assembled high-confidence datasets of missense and non-coding pathogenic variants. After binarizing the probabilities from the four-class output of FIND, we conducted a comprehensive comparison with existing genome-wide tools for predicting the deleteriousness, pathogenicity or functionality of genetic variants. In the task of classifying 19,772 ClinVar pathogenic missense variants and 20,405 gnomAD (*30*) filtered missense variants (*31*), FIND_D demonstrated the best performance, achieving the highest scores in both the AUPR (0.946) and the AUROC (0.940). Tools such as CADD, ncER (*32*), FATHMM-MKL (*33*), ReMM (*34*), and DVAR (*35*) also showed commendable predictive performance (**Fig. 2C** and **Table S5**), suggesting that many current genome-wide predictors can effectively distinguish pathogenic missense variants. Consistent performance of FIND_D was also observed in a separate test dataset of missense variants from ExoVar (*36, 37*), which contains 5,154 pathogenic variants and 3,694 likely benign variants (**Fig. S3A** and **Table S6**). On the other hand, in the task of identifying pathogenic regulatory variants associated with Mendelian diseases, as cataloged by two recent publications (*11, 38*) (44 pathogenic regulatory variants, 58 benign non-coding regulatory variants from ClinVar (*14*)), FIND_D exhibited a considerable advantage (AUPR = 0.906, AUROC = 0.882), surpassing commonly used tools for predicting non-coding pathogenic variants such as CADD, ReMM, and FATHMM-XF (*39*) (**Fig. 2D** and **Table S7**). Additionally, we manually compiled a dataset of 43 high-confidence pathogenic regulatory variants, supported by functional experimental evidence, alongside same 58 benign non-coding regulatory variants from ClinVar. As expected, FIND_D continues to demonstrate excellent ranking performance (**Fig. S3B** and **Table S8**). These results indicate that the binarized version of FIND_D outperforms other genome-wide predictors in the domain of classical pathogenic variant prediction.

### Improved resolution for identifying favored allele with trait-modulating effect

Distinct from all existing prediction methodologies, a primary objective of the FIND model is to stratify genome-wide variants with different fitness effects. We systematically compiled four independent sample sets corresponding to the biological significance of the four categories in FIND, ensuring these samples had not been encountered in prior training processes. These included: 1) 1,912 fixed/nearly fixed variants with potential positive selection signals, identified by selecting alleles in human ancestor quickly evolved regions (HAQERs) with DAF of 0.95 or higher; 2) 451 favored alleles identified by the DeepFavored method (*40*), exhibiting significant molecular and disease/trait-modulating effects; 3) 14,723 low conservative variants from the LD boundary regions of European (EUR) population as likely neutral variants; and 4) 51,396 pathogenic variants selected from the ClinVar database (*19*) with at least one-star evidence supporting their pathogenic status (see details in **Methods**). Initially, we examined the distribution of PHRED scores for these four independent sample types across various genome-wide predictors, including FIND (**Fig. 3A**). As expected, most existing tools, including our FIND_D, ranked ClinVar pathogenic variants highly. However, they struggled to identify favored alleles with potential trait-modulating effects (alleles that aid adaptation to changing environments and lifestyles but also predispose carriers to specific diseases). Notably, existing tools also failed to effectively distinguish functional alleles from neutral ones. In contrast, FIND_I and FIND_N effectively demonstrated the ability to prioritize and distinguish between favored alleles and those that are evolving neutrally or are highly adaptive, a capability not effectively discerned by other existing tools. Additionally, we observed that FIND_F assigns a high rank to corresponding alleles in HAQERs, which may be undergoing or fixed in the process of positive selection (**Fig. 3A**).

**Fig. 3.**
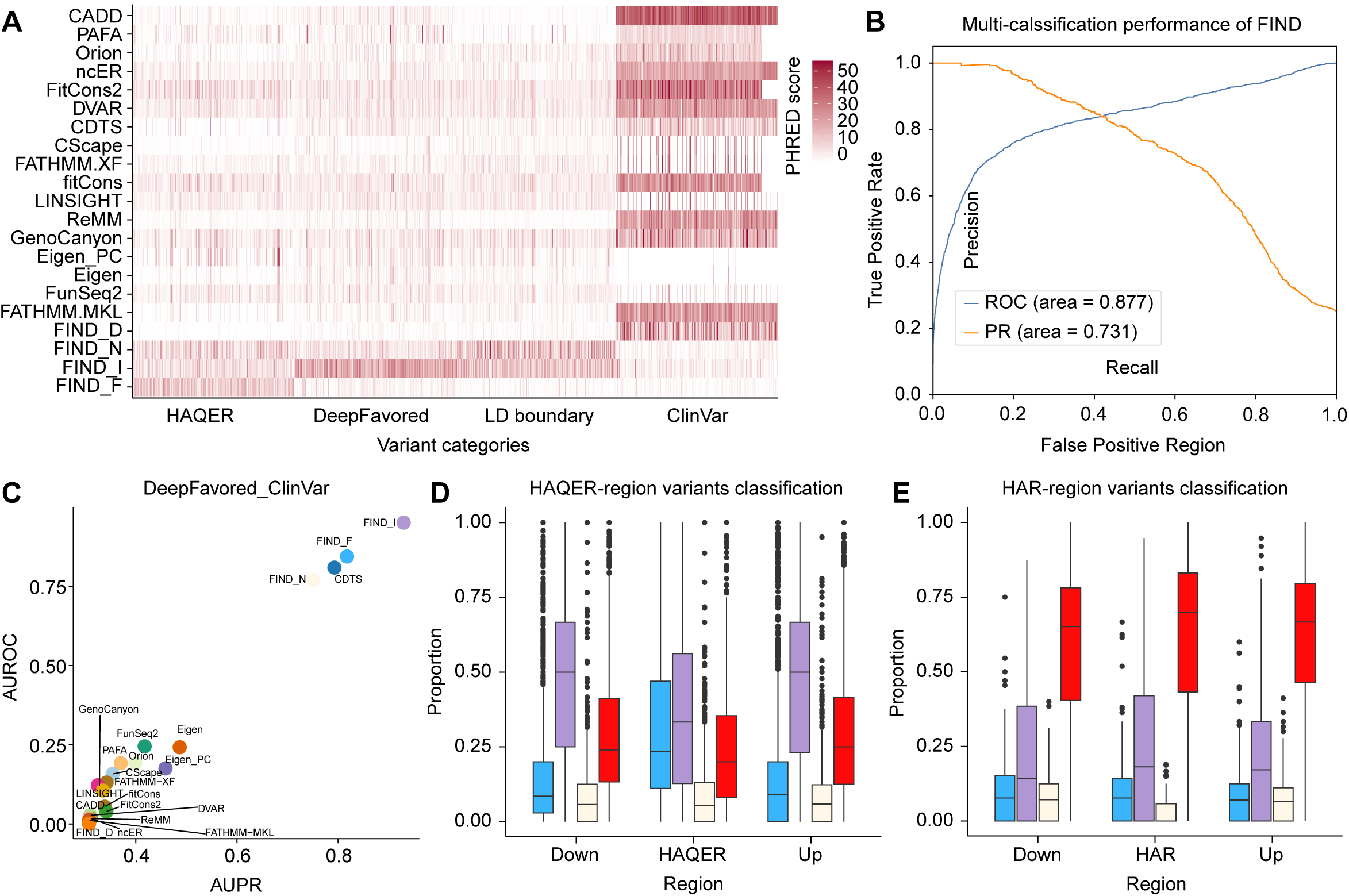
Enhanced resolution in identifying trait-modulating favored alleles. **A**, Heatmap of 17 tools and FIND four categories PHRED score in four-classification test dataset (451 random variants from each filtered data category, see **Methods**). **B**, ROC and AUROC, precision-recall curve (PR) and AUPR for the FIND model evaluation on the first four-classification test dataset. **C**, AUROC and AUPR of 17 tools and FIND prediction from each of four categories in DeepFavored-ClinVar variant dataset. DeepFavored filtered variants as positive sample were labeled 1, while ClinVar filtered pathogenic variants as negative sample were labeled 0. **D**, **E**, HAQER/HAR and flank region variants classification by FIND model. Boxplot was used to show differences in the proportion of these regions that were found to be classified into four categories by FIND.

Using these independent samples for a four-category benchmarking task, we observed that FIND exhibited exceptional performance in this multi-class task (**Fig. 3B**, AUPR=0.731, AUROC=0.877), significantly diverging from existing genome-wide predictors, which primarily recognize and prioritize pathogenic or functional potential. To further explore FIND’s capability in stratifying variant pathogenicity and trait-modulating effects, we employed the binarized version of FIND to distinguish between favored alleles (from DeepFavored data) and pathogenic alleles (from ClinVar data). We found that FIND_I dominate in this task (**Fig. 3C**, AUPR=0.729, AUROC=0.869 **Fig S4A**), while FIND_F and CDTS (*11*) also demonstrate some efficacy in identifying favored alleles (**Table S9**). Interestingly, FIND_D and all other existing genome-wide predictors showed an opposing trend in identifying favored alleles, with FIND_D and CADD being the most extreme. This suggests that nearly all existing tools are designed to identify pathogenic variants and lack the capability to further differentiate favored variants with moderate disease/trait-causal effects. Besides, by substituting two critical variant categories with alternative independent samples (including disease-causal variants identified from the UK Biobank exome-wide association study data (*41*), and likely causal variants from the latest GWAS fine-mapping data), we constructed a second four-classification dataset to test the multi-classification capability of the FIND model (see details in **Methods**). We consistently observed exceptional performance of FIND in such benchmark (**Fig S5A-D** and **Table S10**). Thus, combining the predictive scores of FIND_D and FIND_I could provide a powerful means for effectively subdividing variants based on their fitness impact.

Additionally, we applied the FIND model scores to all known variants in two types of critical human evolutionary regions and their flanking areas to evaluate the model’s specificity in classifying variants in these particular region (*42, 43*). We discovered that, compared to flanking regions, human variations in HAQERs are more likely to be identified with a fixation trend (FIND_F class) and less likely to be recognized as favored (FIND_I class) or pathogenic (FIND_D class) (**Fig. 3D**). This is consistent with the conclusion that both elevated mutation rates and directional positive selection during human evolution drove sequence evolution in HAQERs (*42*). Conversely, in human accelerated regions (HARs), which represent conserved genomic loci with elevated divergence in humans (or regions of the genome with a longer evolutionary time derived from multi-species alignments), the majority of observed human variations are classified as pathogenic (FIND_D class), a proportion higher than in flanking regions (**Fig. 3E**). In these already fixed genomic regions, potentially deleterious variants are emerging. Variations in these HARs predicted as neutral are relatively rare, aligning largely with previous findings (*44*). Together, these results strongly support the uniqueness and accuracy of the FIND model in precisely predicting variant fitness effects.

### Model interpretability and analysis of informative features

Applying the FIND model across the whole human genome resulted in scores (8.6 billion possible base-wise alleles) for four types of variant fitness consequences. To investigate whether FIND’s genome-scale partitioning of variant fitness implicates patterns of recent human adaptation, we first computed the difference of derived allele frequency (*15*) (DDAF) and fixation index (*45*) (FST) for EUR using genotype data from multiple populations within the 1KGP. As expected, we observed a striking decay in DDAF and FST with increasing PHRED scores for FIND_F and FIND_D classes (**Fig. 4A and 4B**), suggesting that the likelihood of variants being predicted as belonging to these classes is higher. This indicates that FIND’s genome-wide predictions for these classes, including preferred alleles that are almost fixed and purified alleles likely pathogenic during human evolution, truly reflect their coordinated allele frequency distributions among human populations. Importantly, since both the training data and applied features were largely derived from EUR genomes and annotations, we found that as the PHRED scores for FIND_I increase, DDAF and FST show a rising trend. This suggests that FIND_I is capable of broadly identifying potential favored alleles in the EUR population that are under selective pressure. In contrast, FIND_N, representing the neutral allele class, did not exhibit any correlation between its PHRED scores and DDAF or FST (**Fig. 4A and 4B**). Together, these results robustly support that the FIND scores can unbiasedly and effectively capture the imprints of recent human adaptation on a genome-wide scale.

**Fig. 4.**
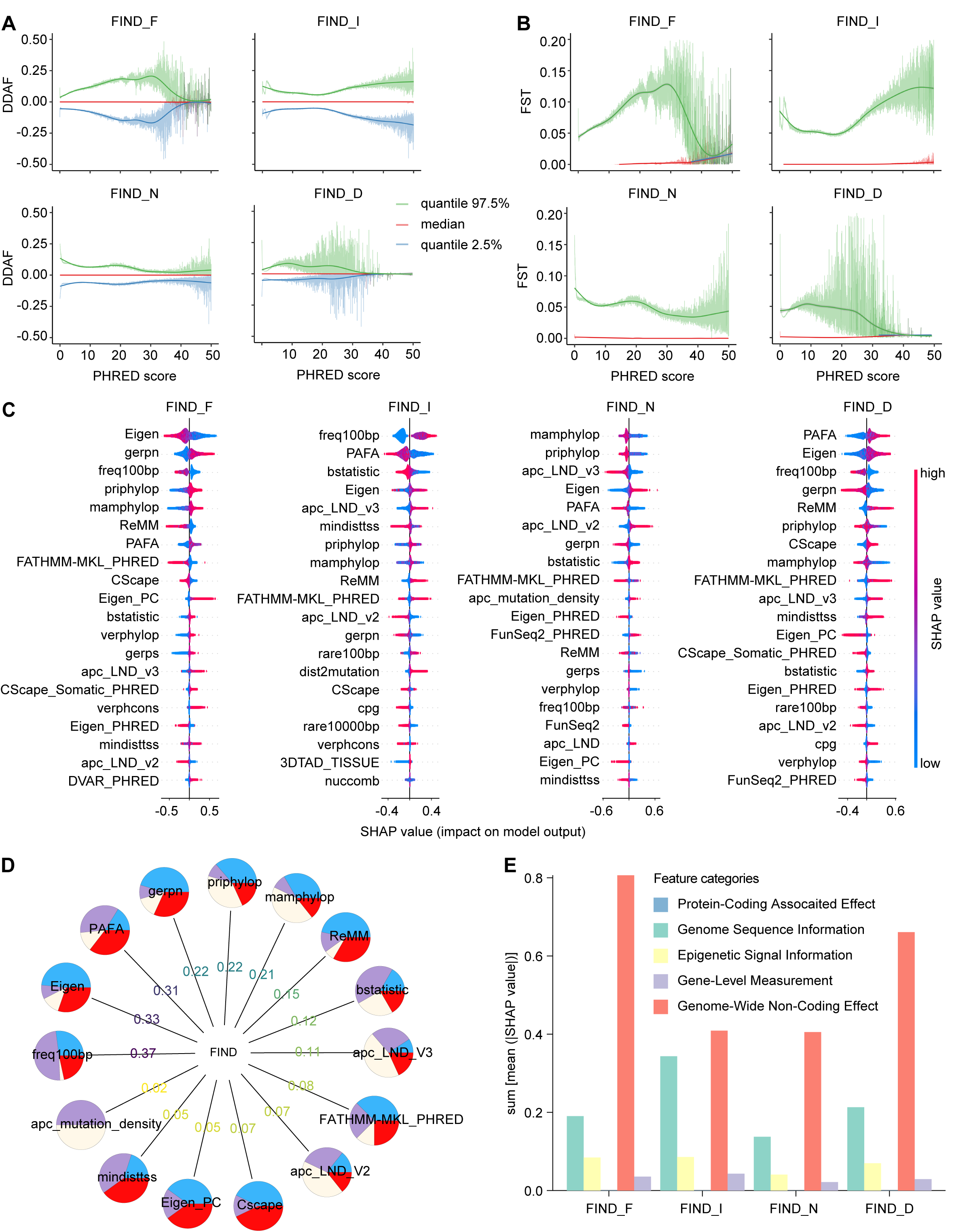
Exploring model interpretability and key feature contributions. **A**, Difference of derived allele frequency (DDAF) score as a function of FIND four-categories PHRED score. The DDAF is the difference in the frequency of derived alleles between the EUR and other four populations (AFR, AMR, EAS, SAS) calculated based on 1KGP data. **B**, Fixation index (FST) score as a function of FIND four-categories PHRED score. The FST is the differentiation of variants between the EUR and other four populations calculated based on 1KGP data. The larger these scores, the more likely the variants have undergone natural selection. For each PHRED score bin (0.01), we selected its 97.5%, 50%(median) and 2.5% quantiles for display. **C**, Summary-plot of top 20 features SHAP values in FIND four-categories prediction (sorted by the mean of the sum of absolute SHAP values of each feature, named mean (|SHAP value|)). LND is short name for local_nucleotide_diversity. **D**, Each pie chart in the outermost layer of the hierarchical relationship diagram represents a distinct feature (union of top 10 features of each FIND category), while the proportion of the pie chart indicates their importance to four different categories variants prediction and the number on the connecting line indicates their importance to FIND model. **E**, Displaying the contribution of five main features categories (the contributions (mean (|SHAP value|)) of features belonging to the same category were summed) to the FIND four categories prediction.

Although the FIND model leverages a deep-learning architecture to extract high-level features from diverse data inputs, it also poses challenges in elucidating the biological relevance underlying its variant fitness classifications. To address this, we utilized SHapley Additive exPlanations (SHAP) (*46*) a game-theoretic approach to interpret the impact of individual features on model outputs at the sample level. Initially, we ranked the top ten most important features for each of the four FIND model categories and aggregated their SHAP values. This cumulative analysis resulted in a union of 15 critical features impacting the FIND model predictions (**Fig. 4D** and **Table S11**). Among these features, sequence attributes such as mutation frequency per 100 base pairs (Freq100bp) and local nucleotide diversity (LND), along with genome-scale pathogenicity scores (including Eigen, PAFA (*47*), and ReMM scores), and evolutionary metrics (such as GERP, priPhyloP, mamPhyloP, and bStatistic (*48*)) were found to influence the FIND model heavily. However, the importance of these features varied significantly across the four model categories (**Fig. 4C**). For example, Freq100bp contributed most significantly to the FIND_I model, indicating its relevance in identifying favored alleles, while Eigen was more influential in the FIND_F model. Additionally, the analysis of the top important features for each model revealed that some features uniquely contribute to specific models. For instance, a novel three-dimensional genomic feature introduced in FIND, 3DTAD_TISSUE (representing the number of tissues where observed variants reside within topologically associating domains (TADs)), had a profound impact. A lower 3DTAD_TISSUE score significantly contributed to predicting non-favored alleles in the FIND_I model, suggesting that variants located within TADs might have more pronounced effects on fitness and human adaptation. SHAP analysis indicated that higher mutation density near observed variants increased the likelihood of being classified as neutral (**Fig. 4C**). Through a comprehensive analysis of five major categories of variant annotations, we observed that the contribution of features to the FIND models was not biased towards specific annotation categories despite different models targeting different variant classes. However, the genome-wide non-coding effects contributed most to the overall model predictions, whereas protein-coding associated effects and gene-level measurements had the least impact, indicating that the non-coding genome plays a significant role in human adaptation (**Fig. 4E**).

### FIND offers fine-grained reclassification for ClinVar variants

Computational approaches have played a crucial role in clinical variant classification tasks (*49, 50*) yet existing tools often categorize variants on a relatively coarse scale, from highly pathogenic to benign. Currently, geneticists and clinicians rely primarily on empirical thresholds or select top-prioritized variants to identify pathogenic variants, leaving them at a disadvantage when dealing with variants of uncertain significance (VUS). Given that the deleteriousness captured by FIND strongly correlates with both molecular functionality and pathogenicity, its multi-class prediction capabilities could provide valuable new insights into this challenge. We utilized FIND to score and categorize all variants listed in the ClinVar 2023-12-30 version (**Fig. 5A** and **Table S12**). Notably, we observed a gradual decrease in the proportion of variants predicted as pathogenic as the ClinVar labels transitioned from pathogenic to benign, with an increase in predictions of trait-modulating variants. Interestingly, even within the ClinVar likely benign variant set, FIND still predicted a significant proportion as pathogenic. This discrepancy could be due to ClinVar’s reliance on genetic, functional, and clinical evidence aiming for higher specificity and adequate penetrance, whereas FIND evaluates the adaptiveness of derived alleles from a broader human evolutionary perspective, capturing subtler deleterious effects and potentially offering a more sensitive assessment of variant pathogenicity. Intriguingly, FIND predicted that 68% of ClinVar benign variants have a trait-modulating effect, suggesting that the majority of non-pathogenic variants catalogued in ClinVar could be functional but might not directly impact human fitness. While FIND demonstrates high sensitivity in identifying pathogenic alleles, leveraging ClinVar’s data on the reclassification of pathogenic variants over the past decade (*51, 52*) could further elucidate FIND’s specificity. We discovered that the PHRED scores assigned by FIND_D to variants initially classified as pathogenic in 2014, but gradually reclassified as non-pathogenic by 2023, were significantly lower compared to other genome-wide variant pathogenicity prediction tools, indirectly representing FIND’s superior control over false-positive rates (**Fig. 5B** and **Table S13**).

**Fig. 5.**
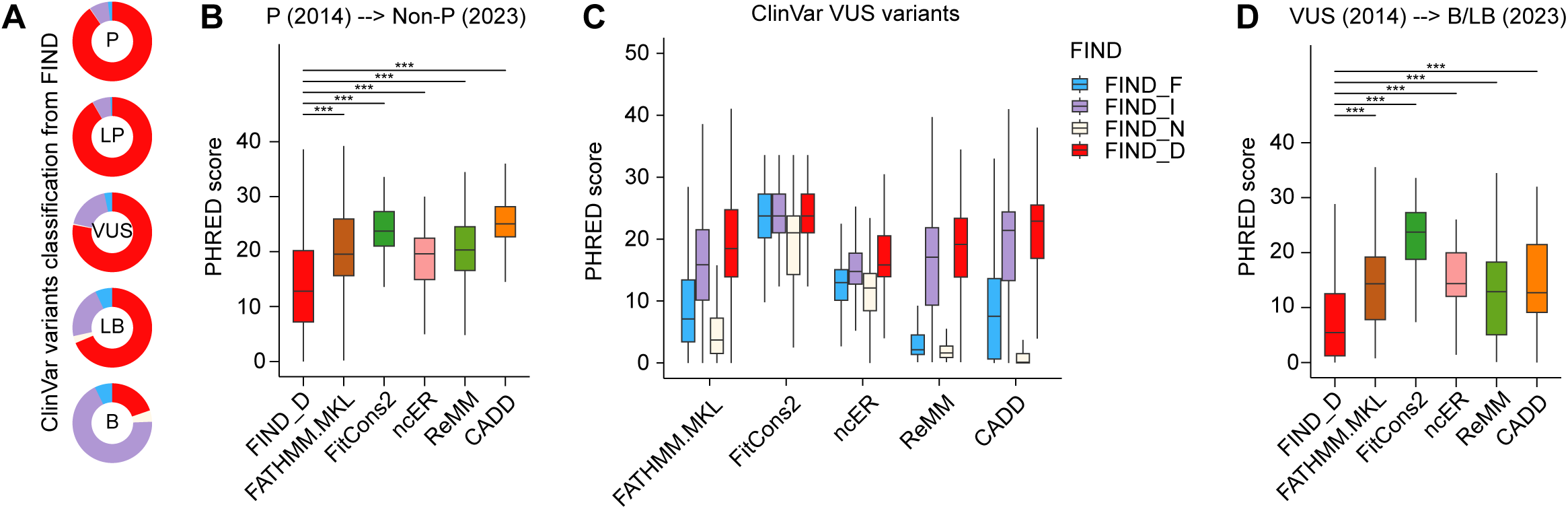
Reclassification of ClinVar variants using FIND model. **A**, Pie chart of FIND classification results for different clinical classifications in ClinVar (P, Pathogenic; LP, Likely Pathogenic; VUS, Variant of Uncertain Significance; LB, Likely Benign; B, Benign). **B**, The distribution of 5 tools and FIND_D PHRED score on reclassification variants from P to B, LB and VUS (namely Non-P), which screened from ClinVar variants 2014 to 2023. *P*-values were computed with Wilcoxon signed-rank test, and asterisk denotes statistically significant differences ***p<0.001. **C**, Boxplot of 5 tools PHRED score distribution base on ClinVar (2023) uncertain significance variant with FIND classification result. **D**, The distribution of 5 tools and FIND_D PHRED score on reclassification variants from VUS to B and LB, which screened from ClinVar variants 2014 to 2023. *P*-values were computed with Wilcoxon signed-rank test, and asterisk denotes statistically significant differences ***p<0.001.

The five-fold increase in variants of uncertain significance (VUSs) deposited in ClinVar from 2020 to 2023 (*53*) poses a critical clinical challenge: accurately categorizing VUSs to enhance the efficacy of genetic diagnostics. In our analysis of 1,078,756 VUSs recorded in ClinVar (2023-12-30 version), 78% were identified by FIND as pathogenic and 18% as favored variants, with the remaining 4% categorized into the other two classes (**Fig. 5A**). Evaluating FIND’s categorization using common genome-wide pathogenic variant prediction scores, we observed a notable discrepancy in the overall PHRED score distribution among tools for FIND-reclassified VUSs. Specifically, excluding FitCons2 (*54*), tools such as FATHMM-MKL, ReMM, CADD, and ncER exhibited distinct median PHRED scores across the four reclassified VUS categories, with FIND_D scoring the highest, followed by FIND_I, and FIND_N showing the lowest scores in most instances (**Fig. 5C** and **Table S14**). This supports the notion that FIND’s classification of ClinVar VUSs holds potential value. However, the distribution of PHRED scores indicates poor discrimination between variant pathogenicity and trait-modulating effects among these tools, complicating the use of specific thresholds to definitively categorize VUSs. This underscores the practical significance of FIND in such tasks. Additionally, we noted that FIND_D showed lower PHRED scores compared to other genome-wide variant pathogenicity prediction tools for variants initially considered VUSs in 2014 but gradually reclassified as benign ones by 2023 (**Fig. 5D** and **Table S15**). In conjunction with the performance noted in the aforementioned ClinVar reclassification records, FIND demonstrates an exquisite capacity to refine the classification of non-pathogenic variants. Together, these results highlight FIND’s potential in effectively distinguishing the clinical consequences of VUSs.

## Discussion

On the whole-genome scale, accurately capturing and defining the contributions and directions of different types of derived alleles to fitness during human evolution and recent adaptation is crucial for understanding the onset of human diseases and the shaping of phenotypes. Traditionally, most computational methodologies have relied on binary dichotomies (e.g., pathogenic versus neutral variants; functional versus non-functional variants) or on sequence conservation for modeling (*55*), which often fails to fully account for complex genetic mechanisms. In this study, we developed a model named FIND, which expands on the traditional binary partitioning approach by categorizing all human genome variants into four nuanced categories, incorporating dimensions of natural selection and the DAF spectrum. This method not only enhances the interpretability of genetic variant impacts but also improves the accuracy in predicting disease-related and adaptive variants. Trained using a deep learning framework, the FIND model focuses on the most critical features at each decision step through its sequential attention mechanisms, thereby not only boosting biological interpretability but also enhancing learning efficiency. Additionally, we systematically integrated and reassessed a wide range of annotation features across different biological domains (*56*). These features provide a rich and informative biological context for the model, enabling effective differentiation between pathogenic and neutral variants across the genome. This refined approach promises significant advancements in our understanding of genetic contributions to human health and disease.

Leveraging two decades of GWAS data accumulation, we have systematically integrated 4,134 full GWAS summary statistics from public repositories. Through statistical fine-mapping techniques, we have pinpointed highly credible disease/trait-causal variants. Unlike pathogenic variants, the majority of these variants exhibit only modest effects on fitness, with many illustrating a trade-off between adaptive evolution and disease susceptibility (*2*). The integration of these likely disease/trait-causal variants from GWAS has propelled the FIND model to unprecedented levels of performance in stratifying variant pathogenicity and trait-modulating effects. Traditional computational strategies generally focus on distinguishing pathogenic/functional variants from non-pathogenic/non-functional ones, thereby failing to adequately identify the full spectrum of derived alleles that are crucial throughout human evolution. However, our GWAS data compilation, predominantly sourced from European ancestry studies, introduces a bias that could potentially restrict the model’s capability to effectively recognize alleles favored by selection in diverse populations. Enhanced genetic research in non-European groups and further studies into recent human adaptation (*57, 58*) are anticipated to address these limitations and enrich our model’s applicability across varied human populations.

The FIND model represents a notable advancement over previous methodologies, enhancing not only the overall performance but also its functionality. First, FIND incorporates novel features that enhance the model’s interpretability. For instance, a range of genome-scale prediction scores, such as non-coding functional scores, demonstrate considerable importance within FIND’s sub-models. Additionally, features related to the 3D genome architecture and epigenetic characteristics have proven to be valuable in predicting trait-modulating variants. These integrated base-wise variant annotations play a crucial role within the FIND model and are also applicable to other variant modeling and prioritization tasks. Moreover, the raw scores of the FIND multi-class model reflect the likelihood of variants, with given annotations, belonging to specific evolutionary selection patterns. This allows for a rapid assessment of the most probable impact of an allele within the current human evolutionary trajectory by evaluating the probability distribution of each allele across different categories. Similar to CADD, by transforming raw scores into PHRED scores, it becomes possible to compare the effects of variants across different types and consequences. This transformation facilitates comparisons among different prediction tools and effectively enhances the prioritization of variants in specific contexts. Lastly, FIND exhibits high sensitivity in classifying clinical pathogenic variants. Leveraging reclassification data from ClinVar over the past decade on pathogenic variants, our findings also suggest that FIND excels in controlling false-positive rates.

While FIND constitutes a significant advancement in variant classification, it is not without its limitations and areas for potential improvement. First, similar to the assumptions made by previous fitness models (*8, 9*), the FIND_D model assesses the "potential of deleteriousness" rather than the "potential of pathogenicity." Unlike limited pathogenicity or functionality metrics, deleteriousness offers a comprehensive, genome-wide assessment free from major biases, capturing the broader impact of variants on phenotype. This makes it a robust measure of variant impact across the genome (*23*). Consequently, interpretations of variant obtaining FIND_D type and its predictive scores require further integration of genetic statistics, phenotypic data, and clinical information to enhance accuracy in identifying disease-causal alleles and in classifying clinical pathogenic variants (*59*). Second, the granularity with which FIND delineates the impact of natural selection and DAF on human fitness could be refined. For example, the contribution of trait-modulating effects to overall fitness within the FIND_I model remains ambiguous, making it challenging to definitively categorize alleles as either adaptive or disease-susceptible. In the FIND_F model, the forces driving fixation might include positively selected alleles, nearby hitchhiking neutral alleles, and even alleles affected by genetic drift. Incorporating finer-grained signatures of positive and negative selection could potentially differentiate these allele types more distinctly (*60, 61*). Furthermore, although FIND provides clear classifications for variants with pathogenic effects and those with trait-modulating effects, the boundaries between these two categories may not be as distinct in reality. Integrating both genome-wide prioritization strategies and multi-class labels could mitigate this limitation.

## Methods

### GWAS full summary statistics curation, integration and normalization

Recent GWASs have amassed a significant number of trait-associated variants that exhibit different fitness in various space-time environment contexts. In this study, we integrated 4,134 GWAS summary statistics based on the European (EUR) population of UKBB from three resources, including Neale Lab UKBB v3 (http://www.nealelab.is/uk-biobank), Gene ATLAS (*62*) and GWAS ATLAS (*63*), and non-UKBB cohorts from several public databases, including GWAS Catalog (*3*), LD Hub (*64*), GRASP (*65*), PhenoScanner (*66*), dbGaP (*67*) and hundreds of summary statistics from websites of consortiums, such as PGC (https://www.med.unc.edu/pgc), MAGIC (https://www.magicinvestigators.org), SSGAC (https://www.thessgac.org), and JENGER (http://jenger.riken.jp/en/). We conducted quality control processing in terms of data recording information, association characteristics information and full summary statistics information: 1) we considered studies that provided clear information, such as population-related information and sample sizes. The mixed populations GWASs or duplicate summary statistics among these resources with less information GWASs were excluded. 2) We considered the trait information within the original GWAS study and manually mapped to Medical Subject Headings (MeSH) terms. 3) Variants with missing summary information that could not be returned were excluded, such as variant coordinate (or rsID), minor allele frequency, effect/non-effect allele, *P*-value, effect size (beta-coefficient) and standard error. In addition, if INFO metric of imputation was available in the raw data, variants with INFO < 0.9 were filtered out.

### GWAS causal variant fine-mapping

Fine-mapping was performed on the curated GWAS data to screen out the causal variants of a trait. We utilized LDetect (*68*) to partition the genetic variants with relatively independent LD blocks and estimated LD information in each LD block by using 1000 Genome Project (1KGP) reference panel (*20*). We employed the FINEMAP (*69*), a widely recognized fine-mapping software, for our analysis. This tool was applied individually to each LD block and associated trait, with assumption that there is a single causal variant within each LD block. We configured FINEMAP using the recommended parameters to ensure optimal performance. Through this approach, we generated posterior probabilities (PP) for each variant, which reflect the likelihood of a variant being the causal entity within the analyzed model. These probabilities were crucial in identifying potential causal variants linked to the traits under study, and were subsequently integrated into a credible set for further analysis.

### Construction of FIND training datasets

The FIND is based on the differences in fitness among genetic variants from different evolutionary processes in modern human genome. Specifically, FIND distinguishes between genetic variants that derived alleles that have exhibited fixed/nearly fixed (labelled **F**), intermediate selection (labelled **I**), neutral (labelled **N**), and deleterious (labelled **D**) patterns. 1) In the course of human evolution, some variants have been retained in the vast majority of the human genome due to selective advantage. These variants are evolutionarily close to being fixed in the human genome, so most of them are harmless. On the other hand, *de novo* variants in the human genome often have uncertain effects and are more likely to lead to serious pathogenic consequences in important genomic regions. By comparing the inferred human-chimpanzee ancestral genome with the human genome, CADD (*8*) screened variants with at least 95% derived allele frequence as fixed variants, and a customized script simulation obtained an equal number of *de novo* variants as deleterious variants. Two randomly selected sets of approximately 550,000 variants each were chosen to represent genetic variants that have been fixed (**F**) during human genome evolution or deleterious (**D**). 2) A large number of phenotypic associated variants have been identified in GWAS studies, most of which are affected by positive or negative selection but not fixed in the human genome, exhibiting different consequences in various spatial and temporal environments. We defined these variants as intermediate selection variants (**I**) and thus integrated the most comprehensive GWAS summary statistics of EUR to date and uniformly identifies credible sets of potential causal variants by using a consistent approach for fine-mapping. We filtered those variants using the following criteria: a credible set confidence interval (≥ 0.9), FINEMAP posterior probabilities (PP ≥ 0.1). As a result, we obtained around 550,000 variants in the human genome. 3) A recent study (*70*) found that high recombination regions are considered to be more efficient in under expressing harmful variants, which can be neutral variants (**N**). They defined these regions as short sequences (≤ 15kb) with a recombination rate higher than 5 cM/Mb in the genome. We used genetic map (*71*) to identify recombination hotspot regions across the whole genome (**Table S16**). Subsequently, approximately 300,000 genetic recombination hotspot variants, independent of phenotype or neutral, were obtained after screening for variants that carried an ancestral allele and had low conservation scores (priPhylop < 0.3, mamPhylop < 0.5, and verPhylop < 0.5) from 1KGP data. We annotated variants consequence by using Ensemble Variant Effect Predictor (VEP) (*72*).

### Feature integration and engineering

We extensively collected and compiled 289 functional genomic and genetic features at the whole-genome level, which are mainly divide into five groups. 1) Genome Sequence Information: Genomic sequences contain a multitude of potential functional information, which are inferred from the sequence statistics of the genome, including GC content, the count of variants within a certain genome window from CADD, the first principal component value of the standardized scores of the Genome sequence information subcategories from FAVOR (*73*) and other statistic data from UCSC project (*74*). 2) Epigenetic Signal Information: Epigenetic markers in the genome are key factors that regulate gene or protein expression, thereby affecting phenotype. We extracted the log10 value score of the epigenetic *P*-value of genetic markers for each variant site from the EpiMap (*18*), Epigenomics Roadmap (*21*), promoter capture Hi-C, and UCSC projects, and used them as the epigenetic annotation feature score for each site. These features typically have continuous non-negative values. 3) Gene-Level Measurement: To augment the putative regulatory effects of non-coding variants. We employed the relevant information of genes closest to the variants as training features. These training features include scores of gene dosage sensitivity, haploinsufficiency, and gene loss of function score (pLI) from a cross-disorder dosage sensitivity map (*75*), FANTOM (*76*), and gene expression levels sourced from GTEx (*77*); In addition to genomic functional information, we utilized various tools developed in recent years to predict the functional, conservative, and pathogenic aspects of variants, obtaining calculated scores from these methods as annotation features. The results of these tools explain the possible functional consequences of variants from different perspectives. 4) Protein-Coding Associated Effect: Predictive scores for variants in the focal genome coding regions obtained from dbNSFP (*22*) database, such as MVP (*78*), MPC (*79*) and MetaSVM (*80*), as well as predictive scores related to splicing variants, such as spliceai (*81*), mmsplice (*82*) and other from CADD. 5) Genome-Wide Non-Coding Effect: Inferred genome conservation scores, such as PhyloP (*23*), PhastCons (*24*), GERP (*25*); variants predictive scores obtained from regBase (*14*) and their corresponding PHRED scores, such as Eigen (*10*), GenoCanyon (*83*), ReMM (*34*). A completed list of features along with their description and accession links can be found in **Table S1**.

All variant annotation data were stored in a tabular format file, which, after compression, exceeds 1.2 TB. We used VarNote (*84*) and custom script to obtain annotations for single nucleotide variants (SNVs). From the annotations described above, some columns that were not useful for model training were excluded. To fit models, we imputed missing values in genome-wide measures by the genome average obtained from the annotation data or setting missing values to 0 where appropriate (**Table S1**). Features from CADD that expand due to the treatment of categorical annotations. There are a total of 906 features after feature engineering and subsequently utilized for model training. All input features and the model were mapped to the human reference build hg19, although annotations for the hg38 version will also be provided through version conversion. Pearson correlation test and hierarchical clustering were used to evaluate the relationships of features upon training datasets.

### FIND model and evaluation

We utilized the TabNet (*26*) model to classify four genetic variant categories in our training set. TabNet model enables the processing of high-dimensional, sparse datasets while maintaining high accuracy and interpretability, making it ideal for classification tasks. The ten-fold cross-validation was conducted on the training set to evaluate the model’s classification performance. For the final training, we used the entire training set and set the following hyperparameters: the width of the decision prediction layer and attention embedding for each mask were 64, the architecture consisted of 5 steps, the learning rate was set to 0.02, and the coefficient for feature reusage in the masks was 1.5. At each step, there were two independent gate linear units and shared linear units. We also trained other machine learning algorithms, including AdaBoost and logistic regression, to compare with TabNet algorithm. The model’s performance was measured using the area under the receiver characteristic operator curve (AUROC) and the area under the precision-recall curve (AUPR), and micro score was calculated because of the four-classification task. We annotated about 8.6 billion variants across the genome with 289 features, and feature processing was carried out. Then, we calculated the FIND four categories probability scores and the corresponding PHRED-scaled (-10*log10(rank/total)) score for each variant. To analyze the distribution of PHRED scores across all possible SNVs specific functional consequences, we first annotated the consequences of all 8.6 billion variants using the VEP and counted the proportion of variant consequences within intervals with consecutive FIND PHRED scores of 1. Subsequently, we standardized the proportion of variant consequences within each scoring interval according to the total number of variants observed in each consequence category.

### Benchmarks on independent testing datasets

We used several datasets extracted from the literature and public databases to look at the performance of the FIND. For the missense variants testing dataset, 1) we first collected the dataset from an state-of-the-art coding variants tool, ESM1b (*31*), which contains 19,772 pathogenic variants from ClinVar (*19*) and 20,405 benign variants from gnomAD (*30*). Besides, 2) After removing variants that could not be annotated to CADD origin features, 5,154 pathogenic variants and 3,694 nearly nonpathogenic variants from ExoVar (*36, 37*) were used as another testing dataset. For non-coding variants testing dataset, 1) we collected 44 pathogenic regulatory variants from two recent publications (*11, 38*) and 58 non-coding benign variants sampling from ClinVar as the first testing dataset (*14*). 2) We also manually collated 43 rare pathogenic regulatory variants supported by functional experimental evidence (**Table S17**) as higher confidence positive pathogenic samples, alongside above 58 non-coding benign variants from ClinVar as another testing dataset. As our model is a four-class variation model, while most existing tools and test sets are binary, we calculated the FIND_D prediction results specifically for the deleterious category of variant. This approach allows for a fair comparison with existing tools and ensures compatibility with binary variant test sets.

Furthermore, to assess the performance of the model in a four-classification task, we constructed a separate test dataset comprising independent variants corresponding to different fitness categories of training data. The test dataset consisted of the following four categories: 1) Fixed variants category: Variants located in the HAQER (*42*) region with a derived allele frequency greater than or equal to 0.95 were screened from the 1KGP dataset, resulting in 1,912 variants that are nearly fixed in the human genome. 2) Intermedia selection variants category: For this category, we collected test datasets from two sources. Firstly, we employed the DeepFavored-filtered variants (*40*), which utilizes the integration of multiple statistical scores to detect signals of positive selection effectively. Specifically, 451 variants from the CEU population, identified by DeepFavored as advantageous sites exhibiting positive selection, were selected. Additionally, we extracted 92,175 variants from the most recent GWAS Catalog that showed significant phenotypic associations (*P*-value ≤ 5e-8), which were not included in the trained dataset. 3) Neutral Variants Category: To assemble the test set comprising likely neutral variants, we identified 14,723 variants located within 500bp upstream and downstream of the LD boundary (*68*) of the EUR population. These variants were selected based on their low conservation scores (priPhylop ≤ 0, mamPhylop ≤ 0, and verPhylop ≤ 0) from the 1KG. 4) Deleteriousness variants category: We selected two datasets corresponding to newly identified deleterious variants in the genome. Firstly, we utilized the ClinVar dataset (2023-12-30) that included 51,396 variants labeled as pathogenic with at least one-star evidence. Additionally, we obtained 446 SNVs with significant disease associations from the UK Biobank whole exome association analysis study (ExWAS) (*41*) data. All datasets mentioned above were carefully filtered to remove any duplicated variants in the training dataset. From the HAQER-filtered variants, DeepFavored variants, LD boundary variants, and ClinVar pathogenic variants, we randomly selected 451 variants to construct the first four-class test dataset. Similarly, we assembled the second four-class test dataset using 446 randomly selected variants from the HAQER-filtered variants, GWAS Catalog variants, LD boundary variants, and ExWAS pathogenic variants. Each dataset was sampled 10 times to ensure the consistency of the results.

### Human evolution analyses

To explore the performance of the FIND model in variant fitness, we evaluated it in two ways. 1) Human accelerated regions (HARs) can be identified by comparing genomes of multiple species, which represent conserved genomic loci with elevated divergence in humans, and also reflect the evolution of the human genome. A recent study identified 2,737 HARs of the human genome by comparing the human genome and genomes of multiple species. These regions are faster to diverge in the human genome, and the variation in these regions may also reflect the uniqueness of our model construction. Besides, 1,550 HAQERs were identified by comparing the inferred human-chimpanzee ancestral genome with the human genome. The evolutionary process of HAQER is shorter than that of HAR. We screened all SNV variants in HAR and HAQERs from 1KGP, as well as their upstream and downstream equal length regions, respectively. Regions with less than 10 variants were removed, and then these variants were annotated, and predicted. After the HARs and HAQERs were screened, 226 and 1,314 regions were left for downstream analysis. 2) DDAF (*15*) and FST (*45*) can reflect the evolutionary differences between different populations to a certain extent. Based on the EUR versus the other four major populations (African: AFR, Ad Mixed American: AMR, East Asia: EAS, South Asia: SAS) of 1KGP data, we calculated the DDAF and FST scores reflecting the selection effect of EUR, followed by 97.5%, 50%, and 2.5% quantiles of the selected scores calculated by FIND PHRED-scale score in 0.01 bin. Given that selected values were predominantly absent or lacking results across the genome, variants devoid of relevant score annotations were excluded from subsequent analyses.

### Feature contribution analyses

Feature contribution was assessed utilizing the Shapley Additive Explanations (*46*) (SHAP) technique employing a permutation explainer. For each feature, we computed the mean sum of the absolute SHAP values (mean (|SHAP value|)) across various classifications within the training data set to capture the feature’s contribution to distinct model classifications. Subsequently, we aggregated the contributions of diverse feature categories, as categorized beforehand into five distinct groups, to determine the overall contribution within each category.

### ClinVar variant reclassification

In 2015, the American College of Medical Genetics and Genomics (ACMG) and the Association for Molecular Pathology (AMP) collaborated to establish comprehensive guidelines to promote uniformity in clinical laboratories (*52*). So, we conducted screenings of the ClinVar dataset at two specific time points (2014-04-01, 2023-12-30). A total of 30,158 and 1,941,252 SNVs were examined in each respective dataset. The classification of these variants was based on specific tags, including pathogenic (P), likely pathogenic (LP), variants of uncertain significance (VUS), likely benign (LB), or benign (B), and variants with at least one-star validation level and non-conflicting variant evidence. We applied the FIND classification prediction method to the 2023 ClinVar variant dataset to calculate the proportion of each category. For tools that use ClinVar variants as the training dataset, the ClinVar classification optimization changes will cause confusion in using these tools. We first screened variants that were converted from P in 2014 to VUS, B, and LB (named Non-P) in 2023, annotated FIND_D and scores of some key tools. Subsequently, we focused on variants that were classified as VUS in ClinVar, the scores of these tools for annotation of variants classified as VUS in ClinVar (2023) were compared with FIND categories. Moreover, the difference of FIND_D and the scores of these tools when variants assigned to VUS categories in ClinVar (2014) change to B/LB in 2023.

## Acknowledgements

The work was supported by the following grants: the National Natural Science Foundation of China (32270717 and 32070675 to M.J.L.), and Technology National Key Research and Development Program (2020YFC2006402 to W.H.Z.).

## Author contributions

M.J.L., W.H.Z. designed the studies and wrote the manuscript. X.T.F., D.D.H. analyzed data and wrote the manuscript. X.R.D., X.F.Y., and J.H.W. evaluated the computational tool and reviewed the manuscript. S.J.Z., X.B.D., X.Q.G., P.C.S., and M.X.L. contributed scientific expertise and reviewed the manuscript. All the authors read and approved the manuscript.

## Competing interests

The authors declare no competing interests.

## Data availability

The annotation information data, FIND prediction raw scores, and relative PHRED scores can be queried from our variant annotation portal (http://www.mulinlab.org/vportal).

## Notes

http://www.mulinlab.org/vportal

### Competing Interest Statement

The authors have declared no competing interest.

